# Cognitive Deficits and Altered Functional Brain Network Organization in Pediatric Brain Tumor Patients

**DOI:** 10.1101/2020.04.22.055459

**Authors:** Hari Anandarajah, Benjamin A. Seitzman, Alana McMichael, Ally Dworetsky, Rebecca S. Coalson, Catherine Jiang, Hongjie Gu, Dennis L. Barbour, Bradley L. Schlaggar, David D. Limbrick, Joshua B. Rubin, Joshua S. Shimony, Stephanie M. Perkins

## Abstract

Pediatric brain tumor survivors experience significant cognitive sequelae from their diagnosis and treatment. The exact mechanisms of cognitive injury are poorly understood, and validated predictors of long-term cognitive outcome are lacking. Large-scale, distributed brain systems provide a window into brain organization and function that may yield insight into these mechanisms and outcomes.

Here, we evaluated functional network architecture, cognitive performance, and brain-behavior relationships in pediatric brain tumor patients. Patients ages 4-18 years old with diagnosis of a brain tumor underwent awake resting state fMRI during regularly scheduled clinical visits and were tested with the NIH Toolbox Cognition Battery. We observed that functional network organization was significantly altered in patients compared to age- and sex-matched healthy controls, with the integrity of the dorsal attention network particularly affected. Moreover, patients demonstrated significant impairments in multiple domains of cognitive performance, including attention. Finally, a significant amount of variance of age-adjusted total composite scores from the Toolbox was explained by changes in segregation between the dorsal attention and default mode networks.

Our results suggest that changes in functional network organization may provide insight into long-term changes in cognitive function in pediatric brain tumor patients.

## 1. Introduction

Primary brain tumors are the most common solid tumors in childhood. With advances in treatment options, overall survival for pediatric brain tumors has significantly improved and the majority of patients become long-term survivors (Gajjar et al. 1997; Packer et al. 2006; Merchant et al. 2009). The presence of a Central Nervous System tumor increases the risk of cognitive impairment, with 20-50% of patients exhibiting such impairment at the time of diagnosis (Krull et al. 2018). In addition to experiencing cognitive deficits from the presence of the tumor, survivors of childhood brain tumors can develop sequelae from their diagnosis and from tumor directed therapies, such as surgery, chemotherapy, and radiotherapy. The sequelae can include cognitive and behavioral performance deficits, compromised employability, and diminished quality of life (Zeltzer et al. 2009; Gupta and Jalali 2017; King et al. 2017; Krull et al. 2018; Ris et al. 2019). Compared to their siblings, survivors of pediatric brain tumors have a 45 fold increased risk of hearing impairment, 8 fold increased risk of visual impairment, and an 11 fold increased risk of unemployment (Gupta and Jalali 2017). Survivors are also less likely to complete secondary education, live independently, or be married (Schulte et al. 2019).

The causes of long-term sequelae in pediatric brain tumors are likely multifactorial, including the structural presence of a tumor during development, hydrocephalus, effects of surgical interventions (e.g., primary tumor resection and shunt placement), the effects of chemotherapy on the developing brain, and the well-documented effects of cranial radiotherapy in children (Hardy et al. 2018; Krull et al. 2018). Sex may also influence long-term effects for survivors, as female pediatric brain tumor survivors generally demonstrate greater improvement post-treatment than males (Bledsoe et al. 2019). However, the precise mechanisms of these injuries are poorly understood, and validated predictors of cognitive outcomes are lacking. Pediatric brain tumor patients undergo serial magnetic resonance imaging (MRI) during their course of treatment and follow-up. Resting state functional MRI (rsfMRI), which can be acquired during clinical imaging, may be a potential modality to assess changes in functional brain network organization that relate to cognitive performance in pediatric brain tumor patients (Matthews et al. 2006).

rsfMRI measures spontaneous changes in the blood-oxygen-level-dependent (BOLD) signal across the whole brain (Power et al. 2011; Kazumata et al. 2017; Roland et al. 2017; Leuthardt et al. 2018). rsfMRI studies in humans have identified anatomically separate, distributed regions of the brain that are functionally linked via strong correlations in these on-going BOLD signals (van den Heuvel and Hulshoff Pol 2010). Such sets of regions form functional brain networks. There is convergent evidence from multiple human and animal studies for the following networks: somatomotor dorsal, somatomotor ventral, cingulo-opercular, auditory, default mode, parietal memory, visual, frontoparietal, salience, ventral attention, dorsal attention (DAN), and reward (Seitzman et al. 2019).

A strength of rsfMRI is that spontaneous BOLD fluctuations can be acquired without requiring patients to perform a task. This freedom from task performance is advantageous over task-based fMRI for testing in young children and patients with cognitive impairments, who may be unwilling or unable to perform adequately on tasks in the scanner. Importantly, rsfMRI, in tandem with network analytic tools, can be used to map the functional network architecture of the brain. Atypical network organization has consistently been identified in patients with brain disorders, including, but not limited to, Alzheimer Disease, Temporal Lobe Epilepsy, and Attention Deficit Hyperactivity Disorder (Fair et al. 2012; Shah et al. 2018; Hojjati et al. 2019).

Formal cognitive assessment by a neuropsychologist, while important for determining clinical care needs, is an expensive and time-consuming assessment for research studies. Utilization of a computer-based cognitive assessment is more feasible in the research setting. The NIH Toolbox Cognition Battery offers a good assessment of cognition that can easily be incorporated into a variety of clinical settings. The assessment is designed to measure executive function, attention, episodic memory, working memory, language, and processing speed for patients ages 3-85 years, and normative data is provided with adjustments for age, sex, race, and parental level of education (Akshoomoff et al. 2014). In this study, we prospectively obtained rsfMRI during routine clinical scanning and evaluated cognitive performance with the NIH Toolbox Cognition Battery in pediatric brain tumor patients.

## 2. Materials and Methods

### 2.1. Participants

Study subjects included patients seeking care at St. Louis Children’s Hospital who were enrolled after informed consent to prospectively undergo resting state functional MRI (rsfMRI) during regularly scheduled clinical imaging along with NIH Toolbox Cognition Battery testing. Eligibility for study entry was age 4-18 years old with diagnosis of any type of brain tumor. Subjects could be newly diagnosed, currently under treatment, or in the follow-up period after completion of treatment. Exclusion criteria were patients with a life expectancy less than one year or patients who were unable to complete cognitive testing due to non-cooperation or motor/visual deficits. The study was approved by the Washington University Institutional Review Board.

### 2.2. rsfMRI Acquisition

All subjects underwent awake, unsedated imaging as part of their routine clinical care. For the rsfMRI scans, subjects were instructed to lie still with eyes open during a period of rest. No auditory or visual stimuli were presented. Scans were obtained in a clinical setting to reduce the cost and time limitations found with obtaining separate research scans. Scans were acquired at various times, including pre-operatively (1), after surgery alone (1), and after surgery plus other treatment modalities, such as chemotherapy and/or radiotherapy (19).

T1-weighted, T2-weighted, and blood-oxygen level-dependent (BOLD) contrast sensitive images were obtained on a Siemens Trio 3T scanner. For the BOLD images, gradient-echo echo-planar imaging with a repetition time (TR) of 2.07s, an echo time (TE) of 25ms, and a flip angle of 90° was used. Two 7-minute runs were acquired, covering the brain in 36 slices (4mm^3^ isotropic voxels) (Fox et al. 2009; Pizoli et al. 2011).

An extant dataset of typically developing children were used to pair each study subject with an age- and sex-matched control. Control data were obtained on a separate Siemens Trio 3T Scanner with very similar acquisition parameters (TR = 2.5s, TE = 27ms, flip angle of 90°, 4mm^3^ isotropic voxels) (Greene et al. 2014).

### 2.3. Preprocessing

To account for magnetization equilibrium and an auditory evoked response to the beginning of the imaging sequence, the first 12 frames of each BOLD run were discarded (Laumann et al. 2015). Slice timing compensation was applied. Then, all BOLD images (for a single subject) were aligned to the first frame of the first run, 3D-cross realigned, and normalized (to a whole-brain mode of 1000). Afterwards, all images were resampled (3mm^3^ voxels), registered to the high-resolution T1-weighted and T2-weighted anatomical images, and then to the Talairach atlas in a one-step operation (Buckner et al. 2004; Smith et al. 2004; Fonov et al. 2011).

Nuisance regression was carried out with regressors extracted from individually-defined ventricles, white matter, and the global mean signal, as well as the 6 parameters from the motion correction step, all of which were submitted to Volterra expansion (Friston et al. 1996; Fox et al. 2009; Power et al. 2014). Segmentation via FreeSurfer 5.3 was used for these individually defined regions (Fischl et al. 2002).

To reduce the impact of motion artifacts, which is particularly important for measuring brain-behavior correlations, frame censoring was implemented. As previously described, all frames with framewise displacement exceeding 0.1mm after applying a low-pass filter on motion parameters were excluded (Siegel et al. 2017; Fair et al. 2020). Frame segments with fewer than 3 continuous frames were also censored. Each BOLD run was required to have a minimum of 30 low-motion frames, and participants with less than a total of 5 minutes of low-motion rsfMRI data were excluded from further analyses (Pizoli et al. 2011). The Washington University NeuroImaging Laboratory 4dfp Suite (https://4dfp.readthedocs.io/en/latest/) and MATLAB (R2015, Natick, MA) were used for all image processing.

A total of 21 study subjects and 21 age- and sex-matched healthy controls were considered for all further analyses.

### 2.4. Functional Brain Network Analysis

Conventional region of interest (ROI) seed-based functional connectivity analysis was performed on the preprocessed rsfMRI data. Functionally-defined ROIs representing “canonical” functional brain networks and regions of the basal ganglia, thalamus, cerebellum, amygdala, and hippocampus were used (Seitzman et al. 2020). Thus, for each patient and control, a correlation matrix was generated. The matrix was sorted into functional networks (defined *a priori*). This matrix represents functional brain network organization for each subject. All ROIs overlapping with either the tumor or resection cavity were excluded from all analyses (replaced with the NaN operator in MATLAB). For some analyses, individual subject correlation matrices were Fisher-Z transformed and averaged together (study subjects and controls in separate groups). In study subjects, ROIs overlapping with a resection cavity (if existent) were excluded for all analyses.

### 2.5. NIH Toolbox Cognition Battery

All study subjects underwent neurocognitive testing via the NIH Toolbox Cognition Battery. Cognitive testing was performed within 6 months of the rsfMRI. Testing was performed in the hospital or clinic setting in a quiet and private environment. All testing was performed with an iPad as prescribed by the NIH Toolbox standard procedures. Test order was standardized by utilizing the same order of tests for all study subjects: Picture Vocabulary Test, Flanker Inhibitory Control and Attention Test, List Sorting Working Memory Test, Dimensional Change Card Sort Test (executive functioning), Pattern Comparison Test (processing speed), Picture Sequence Memory Test (episodic memory), and Oral Reading Recognition Test.

The age-adjusted scores for each measure were exported and analyzed. Additionally, a total of three composite scores were also analyzed. The Fluid Cognition Composite score includes the fluid ability measures (Flanker, Dimensional Change Card Sort (executive functioning)), Picture Sequence Memory (episodic memory), List Sorting Working Memory, and Pattern Comparison. This composite assesses fluid intelligence, which is thought to reflect the ability to learn new tasks or perform known tasks in new situations. The NIH Toolbox Crystallized Composite score includes the Picture Vocabulary Test and the Oral Reading Recognition Tasks, which measure previously learned tasks. The total composite Score includes all of the administered tests.

### 2.6. Statistical Analyses

#### 2.6.1. Brain Networks

For the rsfMRI data, a previously developed omnibus statistical test for functional networks (correlation matrices) was used (La Rosa et al. 2016). Briefly, the technique performs a non-parametric permutation test to determine whether the weighted correlation matrices are significantly different between groups (in this case, study subjects versus age- and sex-matched controls). Then, a post-hoc permutation test was implemented to determine the specific network blocks of the matrix that were significantly different between groups (Gratton et al. 2019). For each test, 1000 bootstrapped samples were run, and False Discovery Rate (FDR) correction was applied to the post-hoc analysis.

Within-network connectivity, a measure of network integrity, was analyzed for the following networks: somatomotor dorsal, somatomotor ventral, cingulo-opercular, auditory, default mode, parietal memory, visual, frontoparietal, salience, ventral attention, DAN, and reward. Within-network connectivity was compared between study subjects and controls using a two-sample t-test. All correlations within a given network were Fisher-Z transformed and averaged prior to the t-test. Study subjects with within-network connectivity 2 standard deviations below the mean of controls or more were considered abnormal.

#### 2.6.2. Behavior and Brain-Behavior Relationships

The mean age-adjusted scores of the study subjects were compared to a normative sample via a one-sample t-test (significance level of 0.05 after FDR correction). All NIH Toolbox Cognition Battery scores, including the total composite, are normalized to have a mean of 100 with 1 SD at 15 (85/115) and 2 SD at 30 (70/130).

To explore the relationship between functional brain network organization and cognitive performance, multilinear regression was implemented in MATLAB. The total composite age-adjusted score for each study subject was selected as the dependent variable. The predictors matrix of independent variables included mean correlation values from network blocks that were found to be significantly different between groups. Since the DAN showed the largest change in study subjects (see Section 3.1.), only significant network blocks involving the DAN were considered in order to minimize multiple comparisons and to avoid a rank-deficient predictors matrix (due to the small number of subjects). Sex was added to the predictors matrix as a covariate.

## 3. Results

A total of 21 children with a primary brain tumor diagnosis and 21 age- and sex-matched controls were analyzed. All known subject demographics, including sex, age at time of scan, race, and ethnicity are presented in **Table 1**, as well as tumor location and treatment modalities for study subjects. The study subjects’ specific tumor diagnoses are presented in Supplemental Table 1. The median time of rsfMRI acquisition was 20 months after diagnosis (range 0 months - 141 months), with only one acquisition occurring less than 7 months after diagnosis. The mean and median time between acquisition of rsfMRI and the NIH Toolbox was 13 and 0 days, respectively, (range 0-178 days) and exceeded 8 days in only two study subjects.

**Table 1:** Subject Demographics.

| **Control Subjects** | **Female (N=5)** | **Male (N=16)** |
| --- | --- | --- |
| Median age at time of scan (range) | 10.6 (8.4-18.1) | 12.6 (9.5-18.6) |
| Race |  |  |
| - White | 4 | 13 |
| - Black/African American | 0 | 1 |
| - Unknown | 1 | 2 |
| Ethnicity |  |  |
| - Non-Hispanic | 4 | 16 |
| - Hispanic | 1 | 0 |
| **Study Subjects** | **Female (N=5)** | **Male (N=16)** |
| Median age at time of scan (range) | 10.7 (8.3-17.9) | 12.6 (8.9-18.9) |
| Median tumor diagnosis age (range) | 7.4 (5.0-13.7) | 9.8 (1.6-17.9) |
| Race |  |  |
| - White | 4 | 12 |
| - Black/African American | 1 | 4 |
| Ethnicity |  |  |
| - Non-Hispanic | 5 | 15 |
| - Hispanic | 0 | 1 |
| Tumor Location |  |  |
| - Infratentorial | 1 | 10 |
| - Supratentorial | 4 | 6 |
| Specific Tumor Location |  |  |
| - Brainstem | 1 | 2 |
| - Cerebellum | 0 | 8 |
| - Pineal | 1 | 2 |
| - Suprasellar | 2 | 4 |
| - Thalamus | 1 | 0 |
| Treatment |  |  |
| - N/A (Pre-Operative) | 0 | 1 |
| - Surgery | 0 | 1 |
| - Surgery + Chemotherapy/Targeted Agent* | 2 | 3 |
| - Surgery + RT | 3 | 8 |
| - Trimodality | 0 | 3 |

RT = Radiotherapy, Trimodality = Surgery + RT + Chemotherapy, \*Vemurafenib

### 3.1. Functional brain network organization is significantly affected in pediatric brain tumor patients

There was a significant difference (*t* = 12.2, *p* = 4.2×10^−38^) in the mean framewise displacement between the study (0.28 +/− 0.47mm) and control (0.20 +/− 0.34mm) subjects. However, after implementing a strict frame censoring threshold, there was no significant difference (*t* = 0.69, *p* = 0.49) in the number of frames retained between study (259 +/− 76) and control (240 +/− 100) subjects.

Within-network connectivity analysis of 12 cortical functional networks was performed. Compared to controls, study subjects demonstrated average within-network connectivity scores greater than 2 SD below the control subject mean in the cingulo-opercular, default mode, parietal memory, salience, and dorsal attention networks (DAN) at a significantly higher rate than expected (*p* < 0.0001; **Fig 1**). The DAN had the largest within-network connectivity deficit for study subjects compared to controls, with 86% of study subjects having an absolute connectivity score greater than 2 SD below the mean of control subjects.

**Figure 1:**
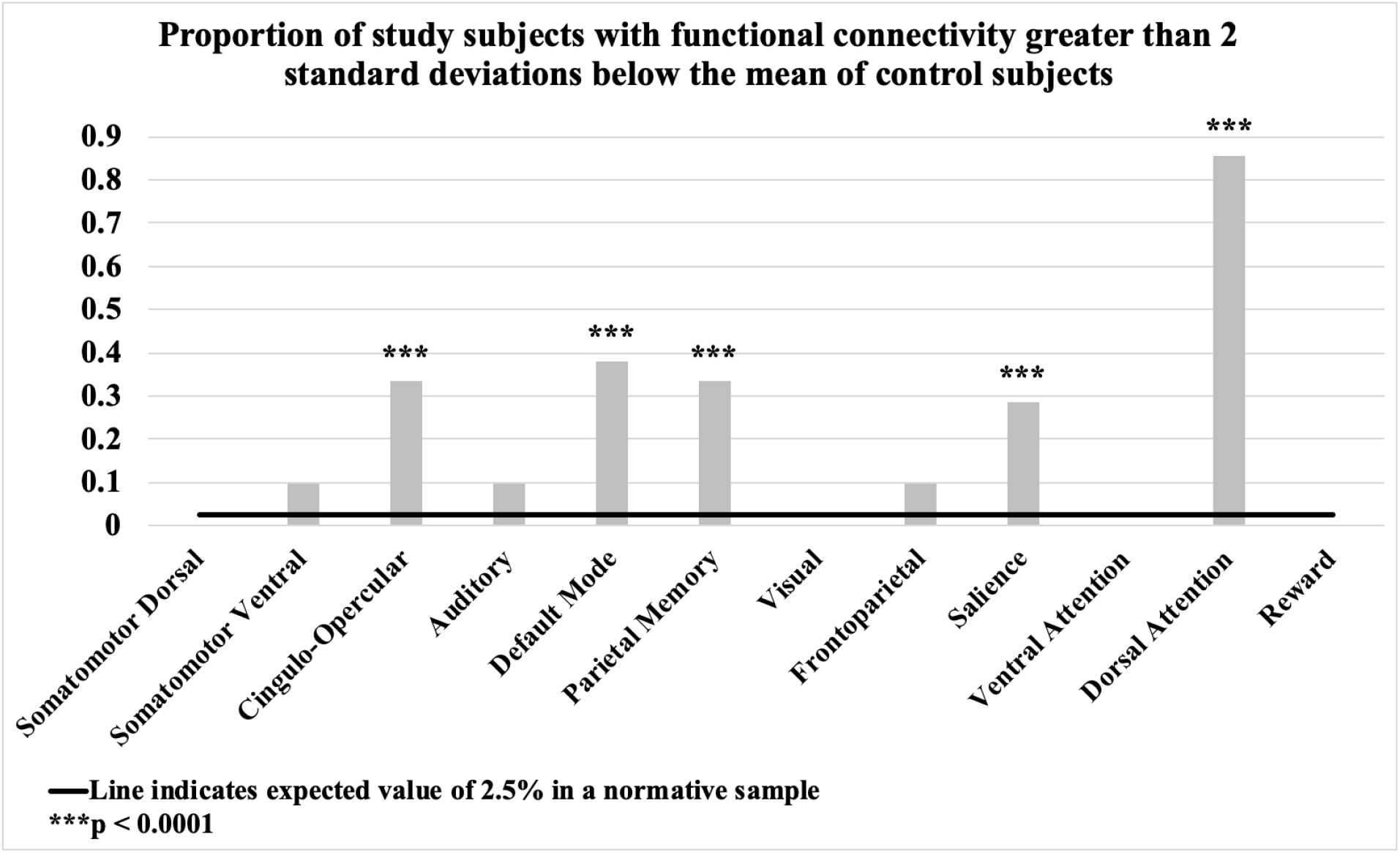
Within-network connectivity in pediatric brain tumor patients. The proportion of patients with functional connectivity greater than 2 SD below the mean of control subjects for 12 selected resting state networks (*** *p* < 0.0001).

There was a significant difference (*p* < 0.001) in overall functional brain network organization between the study and control subjects (**Fig 2A**). Post-hoc analysis revealed that the specific within- and between-network effects (*p* < 0.05, FDR corrected) were within the cingulo-opercular, default mode, parietal memory, salience, DAN, and medial temporal lobe networks, and between the cingulo-opercular and default mode, cingulo-opercular and visual, cingulo-opercular and DAN, cingulo-opercular and reward, auditory and default mode, auditory and visual, auditory and reward, default mode and parietal memory, default mode and visual, default mode and salience, default mode and ventral attention, default mode and DAN, parietal memory and DAN, visual and salience, visual and ventral attention, salience and medial temporal lobe, salience and reward, and medial temporal lobe and reward networks. There were also significant differences within the thalamus, within the cerebellum, between the reward network and thalamus, and between the reward network and basal ganglia (**Fig 2B; Supplemental Table 2**).

**Figure 2:**
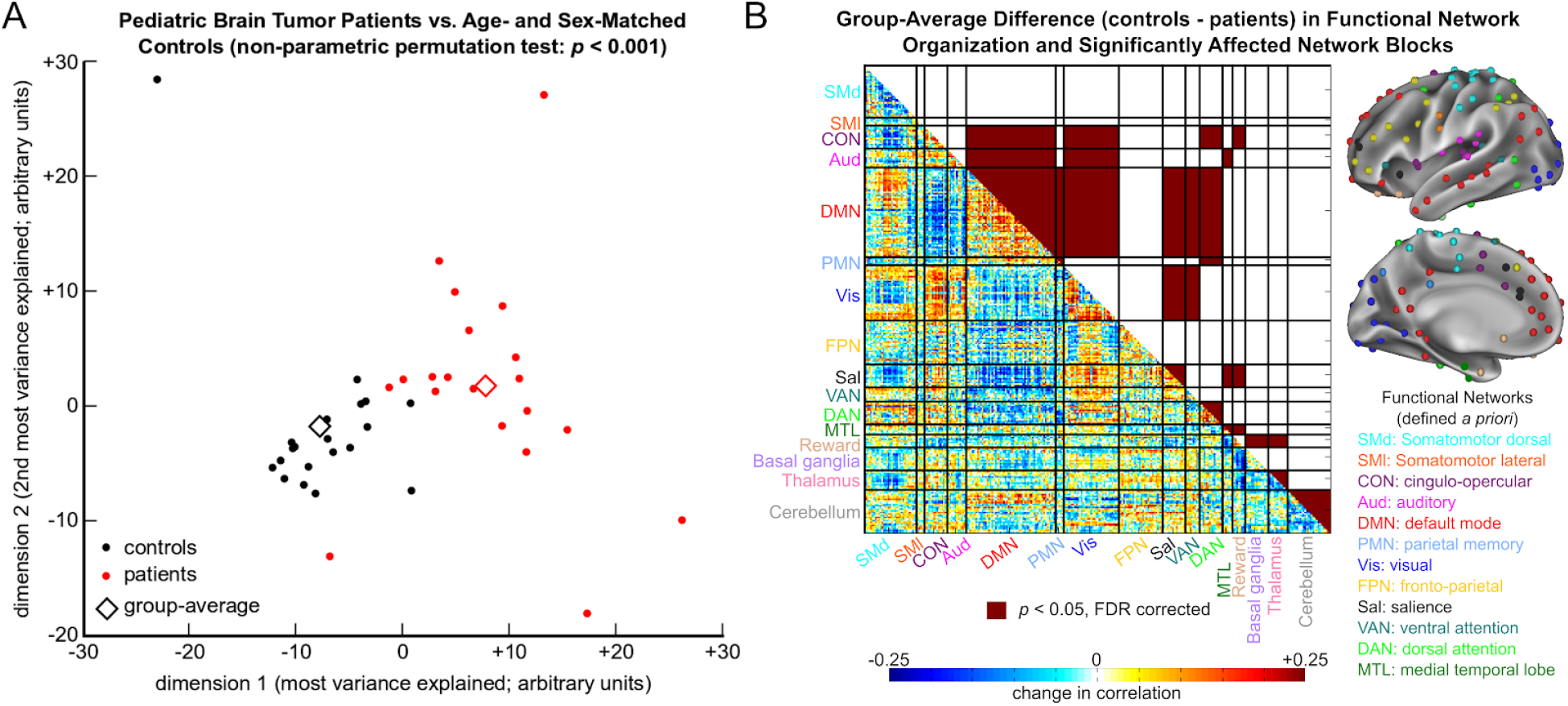
Functional brain network organization in pediatric brain tumor patients. **(A)** The difference between pediatric brain tumor patients (red) and age- and sex-matched controls (black) is displayed. Each filled dot represents the entire correlation matrix (functional network) of a single subject in a multidimensional space, with the group-average for patients and controls represented by diamonds. The two dimensions explaining the most variance in the full dataset are shown. The groups were significantly different (*p* < 0.001) per a non-parametric permutation test designed for correlation matrices. **(B)** The lower triangle of the matrix displays the group-average difference (controls minus patients) in rsfMRI at each region of interest (ROI). Filled blocks in the upper triangle of the matrix indicate significantly different network blocks between groups (*p* < 0.05, FDR corrected). Solid black lines delineate the functional networks (defined *a priori*) and non-cortical structures (the basal ganglia, thalamus, and cerebellum). Cortical ROIs included are displayed on the brain to the right and are color-coded according to their functional network membership.

### 3.2. Cognitive performance is significantly affected in pediatric brain tumor patients and related to brain network organization

Significant deficits (*p* < 0.05, FDR corrected) were found in the domains of attention, cognitive flexibility, and episodic memory when comparing study subjects to the normative mean (**Fig 3A;** **Table 2**). These deficits were reflected in the fluid composite and total composite scores, both of which were significantly different than the normative mean, as well.

**Figure 3:**
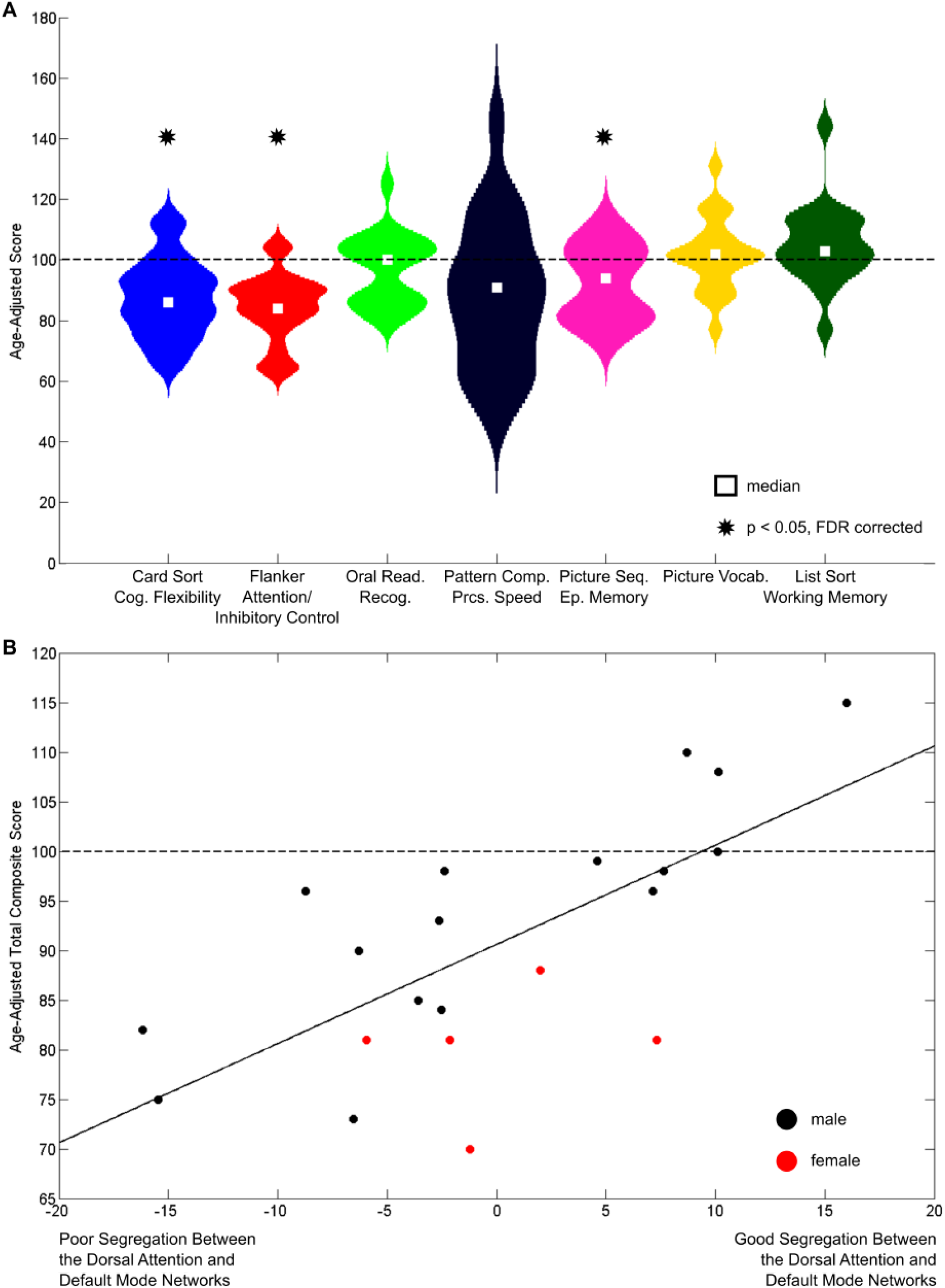
Cognitive performance in pediatric brain tumor patients. **(A)** The violin plots display the distribution of age-adjusted scores for all measures from the NIH Toolbox Cognition Battery obtained from the study subjects. The medians are represented by white boxes, and the normative population average is represented by the dashed line. Significant deficits in cognitive performance were observed in the Dimensional Change Card Sort (cognitive flexibility), Flanker Attention and Inhibitory Control (attention and executive functioning), and Picture Sequence Memory (episodic memory) Tests (*p* < 0.05, FDR-corrected). **(B)** The age-adjusted total composite score from the Toolbox is plotted against the residuals from the multilinear regression model. The sex of each study subject is represented by the color of the marker (red = female), the regression line is represented by the solid line, and the normative population average is represented by the dashed line. A significant amount of variance (R^2^ = 0.52) was explained by sex and the segregation (average connectivity) between the dorsal attention and default mode networks.

**Table 2:**
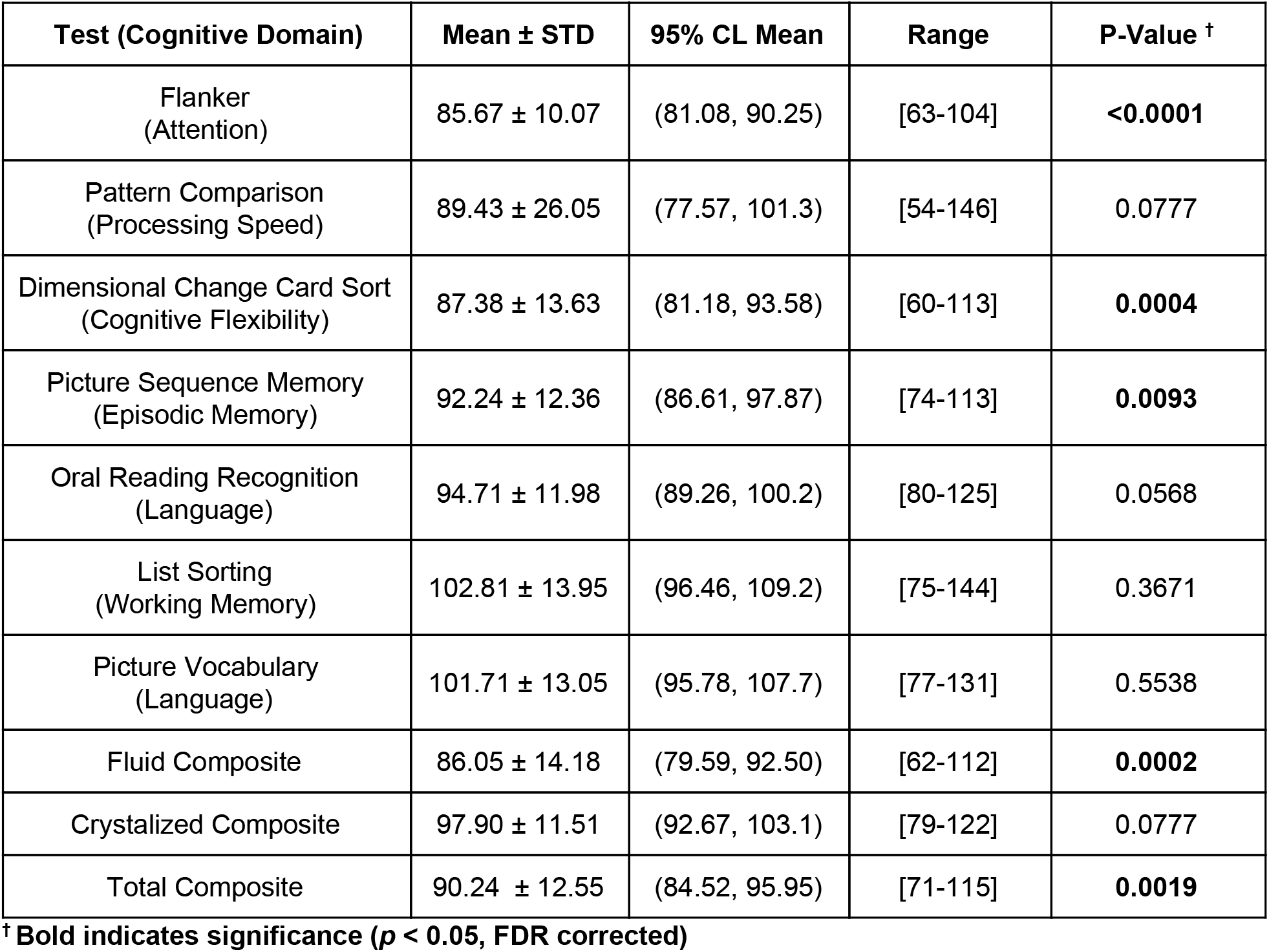
NIH Toolbox Age-Adjusted Scores.

A multilinear regression model explained a significant (F = 3.2, *p* < 0.05) amount of variance (R^2^ = 0.52) of the age-adjusted total composite scores of the study subjects (**Fig 3B**). The only significant predictors were average connectivity between the DAN and default mode network (β = −219.7, 95% CI = [−414.1, −25.3]) and sex (β = +17.7, 95% CI = [6.4, 29.0]). Thus, stronger negative connectivity between the default mode and dorsal attention networks was associated with higher total composite scores from the NIH Toolbox Cognition Battery, whereas weaker negative connectivity and female sex were associated with lower scores.

## 4. Discussion

The present study evaluated rsfMRI and computer-based neurocognitive assessment using the NIH Toolbox Cognition Battery for awake pediatric brain tumor patients. Prior research has revealed that pediatric brain tumor survivors face an increased risk for neurocognitive impairment both pre- and post-treatment (Hardy et al. 2018; Krull et al. 2018).

### 4.1. Brain network architecture and cognitive performance are disrupted in pediatric brain tumor patients

Overall, functional brain network organization was significantly different in study subjects compared to control subjects. Disruptions were observed within and between many networks, particularly within the dorsal attention network (DAN), in study subjects compared to age- and sex-matched controls. Specifically, connectivity strength within the dorsal attention network (DAN) was greater than 2 SD below the mean of the control subjects for 86% of the study subjects. We found corroborating evidence of poorer cognitive performance in the domains of cognitive flexibility, episodic memory, and attention in the study subjects. Furthermore, there was decreased segregation between the DAN and the default mode network in study subjects. This decrease in network segregation, in addition to sex, explained a significant amount of variance in cognitive performance.

### 4.2. The dorsal attention network is particularly affected

The findings of our study revealed significant deficits in the cognitive domain of attention, among others, in the study subjects. Likewise, patients’ functional network architecture was disrupted, particularly in the DAN and its relationships with other networks. The DAN is the brain network responsible for top-down control of goal directed attention (Posner and Petersen 1990; Corbetta and Shulman 2002; Fox et al. 2006). Thus, it is reasonable to observe deficits in both attention-related behavior and attention network organization.

There was a significant decrease in the segregation between the DAN and the default mode network, the two most anticorrelated functional networks of the human brain (Fox et al. 2009). Intriguingly, the extent to which this specific network segregation was intact was significantly related to cognitive performance. This latter finding is concordant with previously published evidence from adult survivors of stroke, which revealed that longitudinal post-stroke recovery of cognitive functions correlates with the return of network segregation, and is also consistent with the importance of maintaining appropriate network segregation in healthy aging (Chan et al. 2018; Siegel et al. 2018). Here, patients with poor segregation between the DAN and default mode network showed worse cognitive decline. These results indicate that the integrity of functional network architecture in pediatric brain tumor patients may be behaviorally meaningful. However, the mechanisms of cognitive decline in these patients are likely multifactorial and require further investigation.

### 4.3. Future directions and limitations

The findings from this study are impactful and highlight the need for further study of rsfMRI in pediatric brain tumor patients. Survivors of childhood brain tumors experience significant neurologic sequelae from their diagnosis and treatment. rsfMRI is a non-invasive method of analyzing brain organization, capitalizing on synchronized neural activity that occurs between functionally related brain regions, in health and disease. The implications of the present data are that rsfMRI may provide valuable information regarding brain organization and its relationship with cognitive performance in pediatric brain tumor patients.

The strengths of this study include its prospective nature, the novel use of rsfMRI in pediatric brain tumor patients, and the temporally correlated assessment of cognition. The results also demonstrate the ability to obtain quality rsfMRI data in the clinical environment, which is of great significance for future use of this technology in pediatric brain tumor patients. Weaknesses of the study include the lack of longitudinal follow-up rsfMRI data and the small number of subjects. Additionally, there is significant heterogeneity in our study subjects with a wide variety of brain tumors, treatment modalities, and ages. Moreover, the data presented here are cross-sectional and, thus, it is difficult to appreciate fully the long-term relationship between functional brain network organization and cognitive performance. Furthermore, deeper assessment of attention in pediatric brain tumor patients (e.g., the Posner task), with the addition of pertinent clinical factors such as tumor type, size, location, and treatment, may be revealing. Lastly, rsfMRI and neurocognitive assessment occur at various times from diagnosis to follow-up. Given the small study size, we were limited in our ability to evaluate known factors affecting cognition, such as hydrocephalus and treatment modalities (e.g., radiotherapy).

## 5. Conclusions

rsfMRI is a non-invasive tool that is able to measure the functional network architecture of the human brain. An important aspect of rsfMRI is that it can be successfully performed in pediatric brain tumor patients during routine clinical imaging. In the present work, patients’ functional network organization was significantly different than controls, particularly in the dorsal attention network. Similarly, patients demonstrated poor performance on measures of goal-directed attention, among others. Finally, the integrity of functional network architecture was related to overall cognitive decline. Future work with a larger cohort of patients will allow for further study of rsfMRI as a potential tool for evaluation of changes to the brain of children with brain tumors and their long-term cognitive sequelae.

## Authors’ Contributions

Experimental Design: D.L.B., B.L.S., D.D.L., J.B.R., S.M.P.

Implementation: H.A., A.M., A.D., R.C., B.L.S., J.S.S., S.M.P.

Analysis: H.A, B.A.S., A.D., C.J., H.G., D.L.B., J.S.S.

Interpretation of Data: H.A., B.A.S., J.B.R., J.S.S., S.M.P.

Writing of Manuscript: All Authors

## Acknowledgements

Research reported in this publication was supported by the Eunice Kennedy Shriver National Institute of Child Health & Human Development of the National Institutes of Health under Award Number U54 HD087011 to the Intellectual and Developmental Disabilities Research Center at Washington University and the Children’s Discovery Institute.

## Supplementary Material

**Supplemental Table 1:** Study Subject Tumor Diagnosis.

| Study Subjects | Female (N=5) | Male (N=16) |
| --- | --- | --- |
| Astrocytoma of Cerebellum | 0 | 1 |
| Craniopharyngioma | 1 | 3 |
| Ependymoma | 0 | 2 |
| Ganglioglioma | 0 | 1 |
| Glioma | 0 | 1 |
| Medulloblastoma | 0 | 3 |
| Pilocytic Astrocytoma | 3 | 2 |
| Pineal Mass | 0 | 1 |
| Pineal Papillary | 1 | 0 |
| Pineoblastoma | 0 | 1 |
| Suprasellar/Third Ventral Mass | 0 | 1 |

**Supplemental Table 2:** Significantly Affected Network Blocks.

| Network Block<br>(stronger in controls) | FDR-corrected *p* | Network Block<br>(stronger in patients) | FDR-corrected *p* |
| --- | --- | --- | --- |
| Between<br>cingulo-opercular and<br>visual | <0.001 | Between<br>cingulo-opercular and<br>default mode | <0.001 |
| Within default mode | <0.001 | Between reward and<br>basal ganglia | <0.001 |
| Within parietal memory | <0.001 | Between default mode<br>and salience | <0.001 |
| Within DAN | <0.001 | Between medial<br>temporal lobe and<br>reward | <0.001 |
| Between visual and<br>salience | <0.001 | Between default mode<br>and DAN | <0.001 |
| Within salience | <0.001 | Between auditory and<br>default mode | <0.001 |
| Between visual and<br>ventral attention | <0.001 | Between parietal<br>memory and DAN | <0.001 |
| Between default and<br>parietal memory | 0.009 | Between default mode<br>and ventral attention | <0.001 |
| Between auditory and<br>visual | 0.022 | Between default mode<br>and visual | 0.009 |
| Between<br>cingulo-opercular and<br>DAN | 0.039 | Between reward and<br>thalamus | 0.016 |
| Within medial temporal<br>lobe | 0.028 | Between<br>cingulo-opercular and<br>reward | 0.022 |
|  |  | Within thalamus | 0.022 |
|  |  | Within cerebellum | 0.035 |
|  |  | Between salience and<br>reward | 0.035 |
|  |  | Between salience and<br>medial temporal lobe | 0.035 |
|  |  | Between auditory and<br>reward | 0.044 |

